# Ytterbium-doped fibre femtosecond laser offers robust design with deep and precise microsurgery

**DOI:** 10.1101/829242

**Authors:** MB Harreguy, V Marfil, CV Gabel, SH Chung, G Haspel

**Author notes:** Equal contribution corresponding authors (SH Chung, G Haspel). **Corresponding authors** Correspondence to Gal Haspel or Samuel H Chung.

## Abstract

Laser microsurgery is a powerful tool for neurobiology used to ablate cells and sever neurites *in-vivo*. We compare a relatively new laser source to two well-established designs. Rare-earth-doped mode-locked fibre laser that produce high power pulsed radiation recently gained popularity for industrial uses. Such systems are manufactured at high standards of robustness and low maintenance requirements typical of solid-state lasers. We demonstrate that an Ytterbium-doped fibre femtosecond laser is comparable in precision to other femtosecond lasers, but with added reliability. It is more precise and can lesion deeper in tissue than a solid-state nanosecond laser. These advantages are not specific to the model system ablated for our demonstration, namely neurites in the nematode *C. elegans*, but are applicable to other systems and transparent tissue where a precise submicron resolution dissection is required.

## Introduction

A focused pulse of laser light can induce accurately localized sub-cellular damage that has been used to approach diverse questions in biology. In the field of neuroscience, laser ablation can target cells (i.e. neurons and glia) by aiming at the nucleus, or sever neurites (i.e. axotomy and dendrotomy), allowing the study of neuronal function and regeneration *in-vivo*. Laser microsurgery platforms often use pulsed lasers at either nanosecond or femtosecond pulse width. Tissue damage caused by either of these lasers is mediated by a non-linear absorption process known as laser-induced plasma formation caused by high strength electric field at the focus of a pulsed laser beam that ionizes molecules to create a bubble of plasma that vaporizes water and tissue^1,2^. The energy intensity at the focal point that is required to achieve laser-induced plasma formation scales with the pulse energy divided by the pulse duration^3^. Hence, the ten-thousand-fold longer nanosecond laser pulse requires higher energy for plasma formation and ablation, Accordingly, pulse energies used for nanosecond pulse ablation are typically tens of mJ, which generate more severe damage beyond the region of laser energy deposition, compared to tens of nJ for femtosecond pulse ablation^4^.

Laser microsurgery was pioneered and has been an important tool to study the development and neurobiology of *Caenorhabditis elegans* since 1980 when Sulston and White used it to study cell-cell interaction in post embryonic stages^5^, and when Chalfie and colleagues ablated neurons to test their necessity for touch sensitivity^6^. Microsurgery of neurites (termed axotomy) was also pioneered in *C. elegans*^7^ where a laser-pumped titanium-sapphire laser was used to cut commissure neurites of motoneurons. These Ti:Sapphire lasers are typically configured to produce near-infrared pulses with a centre wavelength of approximately 800 nm, pulse energies up to 50 nJ, repetition rates of 80 MHz, and pulse duration of 100–200 femtoseconds. Ablation at 80 MHz does not allow complete dissipation of pulse energy ^8^and in many cases an external electro-optic pulse picking device is added to reduce the repetition rate and ablate at 1 kHz. At this lower repetition rates the deposited energy dissipates completely during the intervals between pulses ^9,8,10^improving the surgical resolution. Nanosecond lasers were adapted for axotomy, offering lower cost, and more robust design^6,11^, but with higher energy per pulse. They are typically Nitrogen laser-pumped or diode laser-pumped dye laser typically configured to produce ultra violet (337 nm), blue (450 and 488 nm), or green (532 nm) light with pulse durations of several nanoseconds and pulse energies up to several mJ with pulse repetition rates up to 10kHz.

Axotomy using high and low repetition rate femtosecond pulses, and nanosecond pulses vary in the size of a gap induced in a severed axon and in the extent of damage to surrounding tissues, but it appears that axon regeneration occurs at comparable rates and extents after axotomy^12^. At low repetition rate (1 kHz), the size of damage to tissue and neurite regeneration rates appear to be mostly affected by pulse energy^13^. To improve surgical resolution it is preferable to use laser energies near the damage threshold. For any energy level there is a minimal number of pulses that will initiate ablation but adding pulses does not substantially increase the region damaged ^3,13^.

Nanosecond lasers advantages are their relatively low cost, compact size, low maintenance and lower safety requirements. Femtosecond lasers dissect with higher resolution and less damage to surrounding tissues. The choice of laser system is usually motivated by application and involve a compromise, considering precision, cost, and reliability. Neurites spaced further apart require less surgical resolution while bundled neurites often require submicrometer resolution^10^. Here we compare a new type of laser, an Ytterbium-doped fibre femtosecond-pulse laser, with the two laser types most commonly used for laser axotomy in *C. elegans:* a femtosecond Ti:Sapphire and a nanosecond diode-pumped passively Q-switched solid state laser.

## Results

### Ytterbium-doped fibre femtosecond laser

Rare-earth doped mode-locked fibre laser produces high power pulsed radiation by amplifying seed source radiation through a thin coiled fibre. Specifically Yb2O3-doped fibre lasers emit pulses that are about one hundred to hundreds of femtosecond wide, at the range of 1000-1100 nm, at average power of mW to tens of watts^14^. The basic design is similar to Ti:Sapphire lasers that are very commonly used for multiphoton imaging and axotomy^15,16^. One advantage of Ti:Sapphire laser over Yb-fibre is that the wavelength is tunable, but this advantage is not crucial for ablation applications such as ours. The advantages of Yb-fibre are higher possible power, lower maintenance, smaller footprint, and air cooling. The specific system used here (BlueCut, Menlo Systems GmbH, Germany) includes an internal pulse picker which simplifies set up and lowers the overall cost compared to Ti:Sapphire systems that require an external pulse picker. Our Yb-fibre system can ablate with user-defined repetition rate of single shot to 50 MHz and pulse energies of nJ to μJ. As further described in the Methods, in the Yb-fibre ablation setup the laser beam is sent through a beam expander into a microscope objective that focuses the pulses into a sample. Brightfield and fluorescent light is captured by a sCMOS camera for visualization and targeting (Figure 1a).

**Figure 1.**
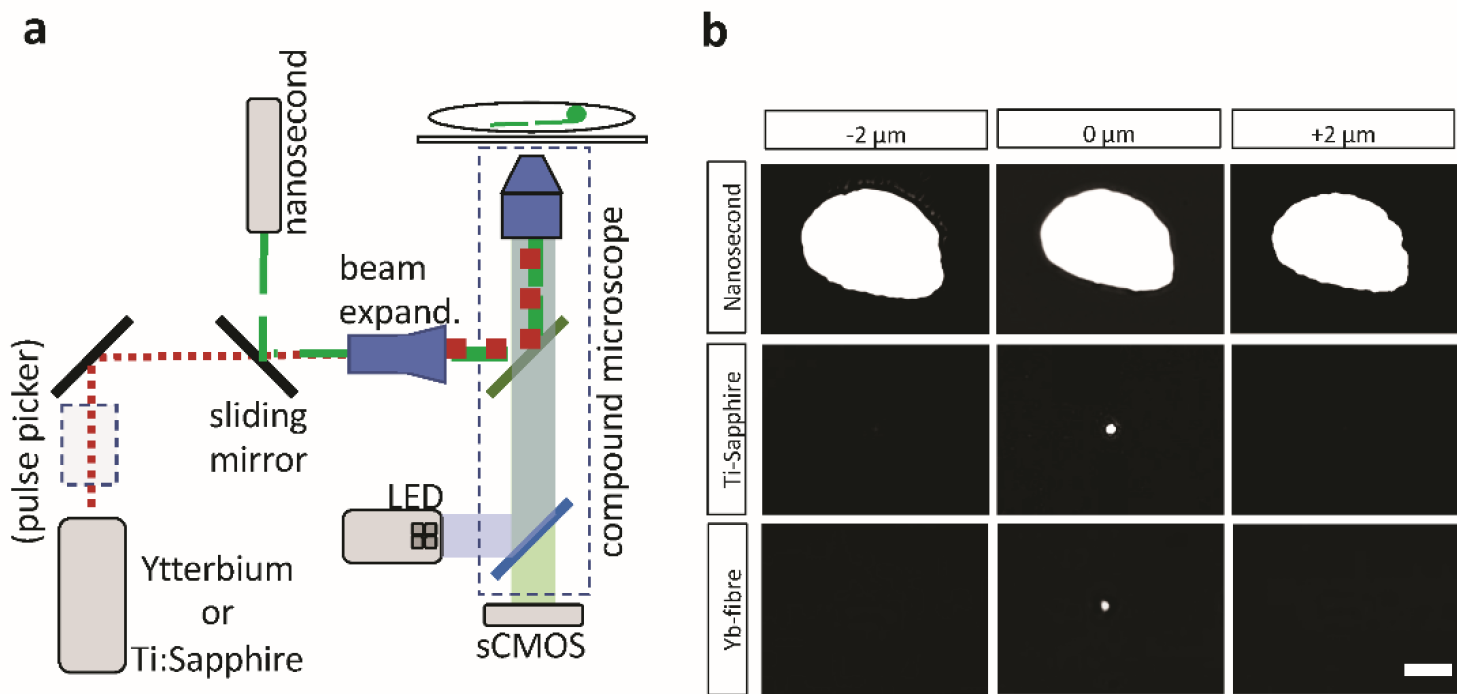
Yb-doped fibre femtosecond laser integrated in epifluorescence microscope produces a small damage spot, comparable to Ti:Sapphire laser. a) Lasers are integrated to an epifluorescence compound microscope through a beam-splitter or dichroic mirror. Note that some lasers require an external pulse picking device. b) Damage spot on a thin layer of black ink is larger when induced with a nanosecond pulse laser (top) than with either of the femtosecond lasers (middle Ti:Sapphire; bottom Yb-fibre). Damage spots are not reduced in size two micrometres above or below the image focal plane for the nanosecond pulse laser, while undetected for the femtosecond pulse lasers. Scale bar = 5 μm.

### Lesion size

Lesion size was evaluated by focusing the laser beams through a coverslip onto a layer of black ink (Figure 1b). The laser was adjusted to the lowest possible power setting that induced axotomy. The Yb-fibre laser and the Ti:Sapphire produced similar damage (p<0.001; diameter 1.2 ± 0.2 μm) at the focal plane and no significant damage 2μm above or below the focal plane. The nanosecond-pulse laser produced a much larger lesion (p=0.43) not only at the focal plane (12.5 ± 0.5 μm) but also 2 μm above or below (13.0 ± 0.4 μm; 14.0 ± 0.4 μm). Even though smaller lesion sizes were reported for the nanosecond laser, that study utilized a very low energy setting (10 pulses at 100Hz) only meant for alignment and not for axotomy^11^.

### Qualitative accuracy test by dendritic bundle ablation in *C. elegans*

A Ti:Sapphire laser system can selectively ablate one sensory dendrite in a tight bundle of twelve dendrites^16^. These dendrites are sensory organs of the twelve amphid neurons located in the nose of *C. elegans*. We replicated this treatment using both the nanosecond pulse laser and the Yb-fibre laser. The Yb-fibre femtosecond laser, similar to the Ti:Sapphire, is capable of ablating only one neurite in the bundle with no damage to surrounding tissue. However, the nanosecond-pulse laser produces a larger injury affecting surrounding tissue and therefore damages more than one dendrite (Figure 2). The damage observed for the nanosecond-pulse laser is more localized than what we previously described in figure 1b, this could be due to the fact that the higher energy might be causing thermal damage to the ink while not vaporizing the tissue. Additionally, the three-dimensional geometry and the ability of the energy to penetrate through the tissue also need to be considered.

**Figure 2.**
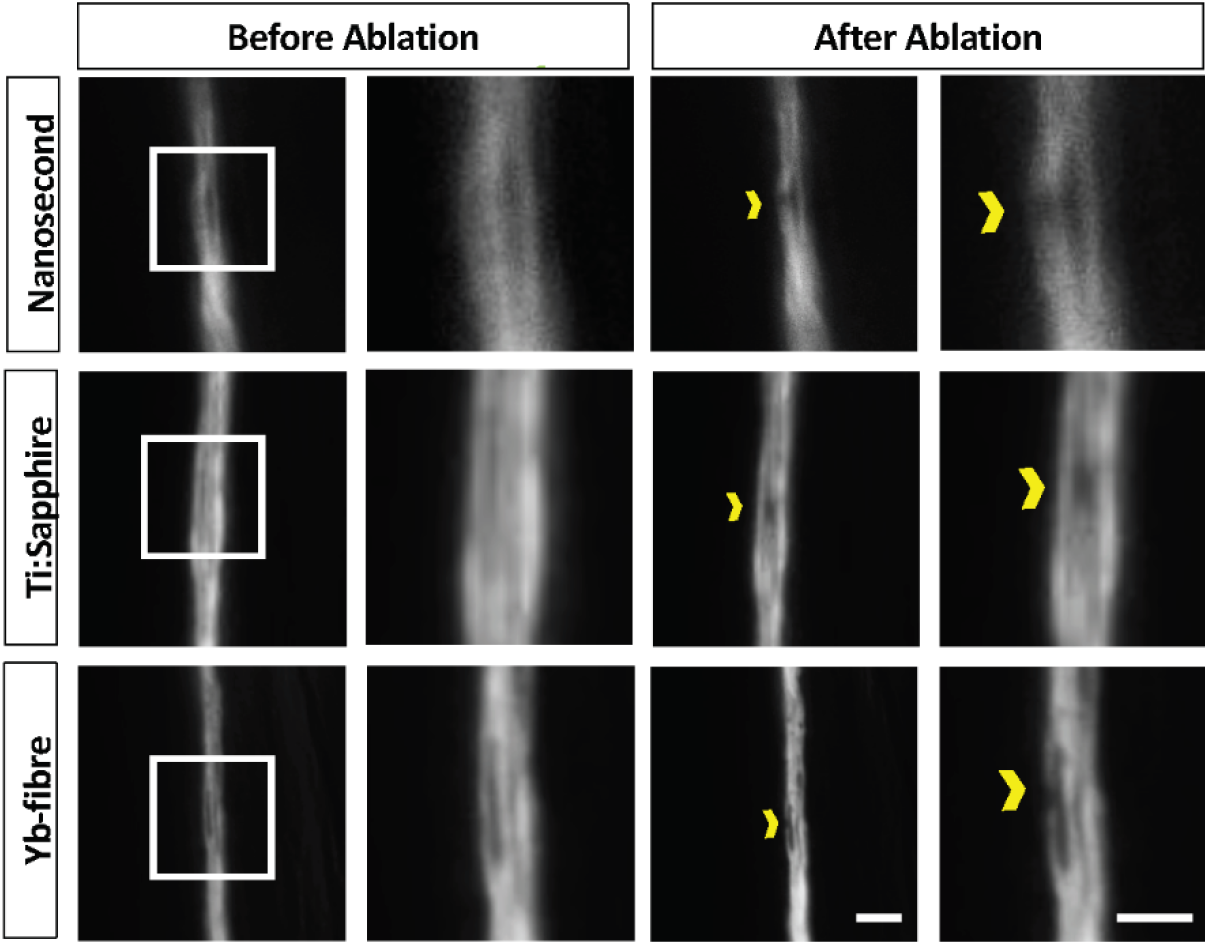
A single *C. elegans* dendrite can be injured without damage to neighbouring dendrites. A single dendrite located in a sensory bundle in the nose of the animals was successfully ablated using the Ti:sapphire and the Yb-fibre lasers without collateral damage to adjacent dendrites. The nanosecond-pulse laser injured more than one dendrite. Scale bar = 2μm.

### *Quantitative regeneration assessment in* C. elegans *motoneurons*

Finally, we assessed how the different laser setups might impact neuronal regeneration in *C. elegans*. Because Ti:Sapphire axotomy has been widely described in the literature^7,17^, we limited our comparison to the Yb-fibre laser and the nanosecond-pulse laser (Figure 3). Animals were immobilized and D-type motoneuron commissures which extend from the ventral to dorsal side of the animal were axotomized (Fig. 3a). The size of the gap (Fig. 3b) between severed ends produced by the nanosecond laser after initial retraction (3.8±1.4 μm) was significantly larger (p=0.02) than the gap produced by the Yb-fibre laser (2.4±0.8 μm).

**Figure 3.**
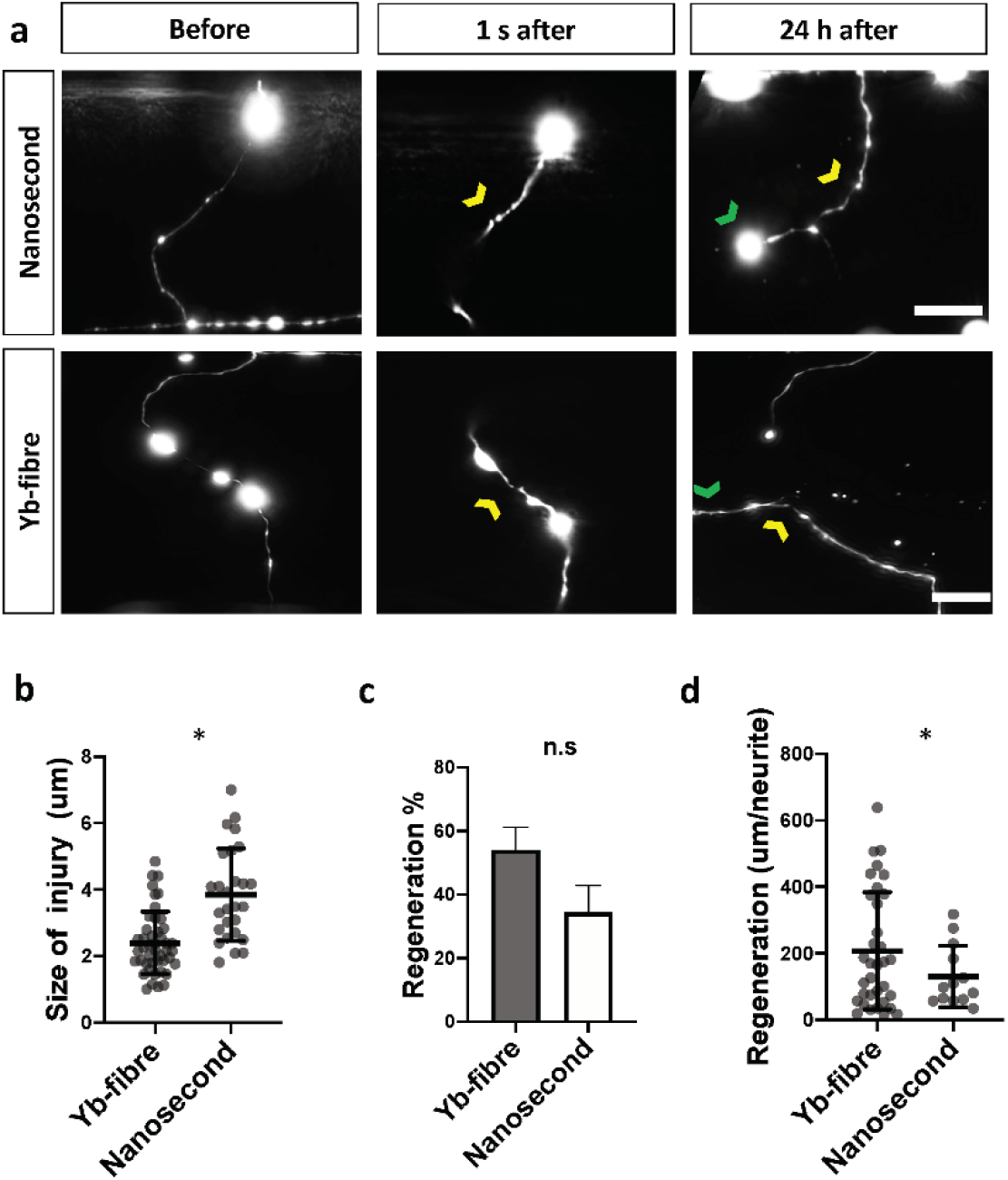
Nanosecond laser induces larger gap at site of injury but injured neurons regenerate at similar rate. a) *C. elegans* motoneuron axotomy. *C*ommissures of D motoneurons in immobilized animals were axotomized, animals were recovered and axon found again next day to asses regeneration. Images taken before, 1 second after and 24 hours after injury. 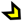 Indicates axotomy site and 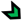 indicates the regenerating branch. b) Mean gap between injured tips was larger after injury with nanosecond laser. c) Percentage of neurons that regenerated at 24 h was the same for the two types of lasers. d) Length of regeneration was slightly larger 24 h after injury with Yb-fibre laser. Images in A for before and at 24 h are maximal projections of a Z stacks while the images at 1 s are each a single frame, leaving some parts of the neuron out of the imaging plane. * p<0.05. Scale bar. = 5 μm

Similar proportion (Fig. 3c; p=0.12) of neurites injured with the Yb-fibre laser regenerated (27 of 50) to those injured with nanosecond pulse laser (11 of 32). However, neurites injured with the Yb-fibre laser extended longer processes (Fig. 3d; 195.8±173.7 μm, n= 50 neurons from 11 animals) than neurites injured with the nanosecond-pulse laser (142.5±102.5 μm, n= 32 neurons from 11 animals; p=0.02).

## Discussion

Yb-doped fibre lasers possess several advantages over more commonly used nanosecond pulse lasers and similar capabilities to Ti:Sapphire lasers. Femtosecond fibre laser systems are designed for the high standards of industrial cutting and welding. Hence, the Yb-fibre laser used here offers integrated power adjustment and pulse picking, controlled by a dedicated software that can pick as few as a single pulse. For these reasons, compared to a Ti:Sapphire system, the use of Ytterbium-doped fibre laser requires much less daily alignment and optimization and less maintenance. It is our experience that a system that was turned off for three weeks could cut axons as soon as it was turned back on. Comparable turnkey laser systems, such as the Spectra-Physics Spirit, are commercially available. However, we found that the BlueCut Yb-fibre laser was the most suitable at the time we initiated this study. The most notable reasons include the integrated pulse picking, the simple incorporation of this unit to an existing microscope, air rather than liquid cooling and the relatively lower cost (including the external pulse-picker required for other systems). In our experience, we never use the full laser power and a unit with about 20% of existing power would have been preferable, for safety and cost reasons.

The ability to produce injury depends on pulse energy and duration. The injury producing plasma is induced by a multiphoton absorption that requires a minimal intensity threshold which is only reached at the focal plane. In the case of neuronal injury with a 1.4 NA objective and a pulse duration of 100 fs, this threshold occurs at a level of energy of 6.5 x10^12^ W/cm^2^ or 4 nJ per pulse^1^. In this study, Yb-fibre and Ti:Sapphire lasers produced comparable pulse width (<400 fs), while the nanosecond-pulse laser produced four orders of magnitude longer pulse (<1.3 ns). Hence, the former induces axotomy with lower energy levels and therefore less collateral damage as longer pulses require more energy to reach the threshold intensity. Yet, regeneration rates were comparable among all three systems.

Further, because near infrared photons penetrate deeper and are less scattered by biological tissue, lower energy levels can be used, and the Yb-fibre laser can axotomize about 100 microns deep into tissue compared to 30-40 microns for nanosecond laser systems^11^. In *C. elegans* this capability permits the ablation of neurons located on the side of the worm farther from the objective.

Here we have described a new laser axotomy system that provides the same high precision capabilities of widely-used Ti:Sapphire lasers but with less maintenance and higher robustness. This capability is not limited to neurons in *C. elegans* and might be useful to address experimental questions in transparent tissue that require an accurate and localized submicrometer-sized lesion up to 100 μm deep with no collateral damage.

## Methods

### Laser platforms

The Yb-fibre system which generates ∼400 fs pulses in the infrared (1030 nm) (BlueCut, Menlo Systems GmbH, Germany) includes an integrated pulse picking unit composed of two acousto-optic modulators (AOMs). A fixed AOM reduces repetition rate from 50 MHz to 1 MHz, a second variable AOM reduces it further as low as 1 kHz. We use the BlueCut Control software provided by Menlo Systems with the laser. The software allows user control of the repetition rate (typically, we use 1 kHz) and external gating of the variable AOM by a transistor-transistor logic (TTL) signal. We use a function generator (model 33210A, Keysight Technologies) to provide the TTL pulse at various lengths (typically 100 ms for 100 pulses at 1 kHz). Reliable picking of single laser pulses can be accomplished through synchronization of the two AOMs in the pulse picker unit.

Alternatively, we pick a single laser pulse with a 1-ms TTL signal, accepting occasional fails that are easy to visually recognize. Users control laser power by choosing a level of 1 to 99 arbitrary units that are scaled by the software. Typically, we use 10-20 units for ablations. We measured 3-65 nJ/pulse at the image plane (PM100D Power and Energy Meter Console with S170C Microscope Slide Power Sensor Thorlabs GmbH) when changing the arbitrary units scale from 4 to 35 (of 99).The beam is directed through a beam expander (10X Achromatic Galilean Beam Expander, AR Coated: 650 - 1050 nm, Thorlabs, USA) and dichroic mirror (750 nm long-pass, Thorlabs, USA).

The diode pumped passively Q-switched solid state system (1Q532-3), Crylas Laser Systems, USA^11^, was integrated via a flip-mounted mirror to the same optical path and microscope objective as the Yb-fibre laser. The only components that had to be replaced were the beam expander (10X Achromatic Galilean Beam Expander, AR Coated: 400 - 650 nm, Thorlabs, USA) and 1:1 beam splitter (Thorlabs, USA) to accommodate the shorter wavelength (532 nm).

The Ti:sapphire femtosecond laser (Mantis PulseSwitch Laser, Coherent, Inc.) generates 100-fs pulses in the near infrared (800nm). We operated the laser at 10 kHz, used 0.25 s exposure for ablations, and propagated the beam through a home-built 10x Galilean expander.

### Ablation parameters

We ablated samples with 13 to 15-nJ (Yb-fibre), 10-nJ (Ti:Sapphire), and 28-μJ (nanosecond) pulse energies. Ablations occurred at 1 kHz, except for *in vivo* Yb-fibre ablations were carried out with 10 kHz repetition rate. We used 100 pulses in all cases except ink ablations with the nanosecond laser, which utilized 5 pulses. We focused pulses with an Olympus UAPO 40X, 1.35 NA oil immersion objective for ink and bundle ablations and an Olympus 100×, 1.4 NA oil immersion objective for motoneuron ablations. Where possible, laser power and pulse number were set to the minimum setting in which damaged could be observed.

### Measurement of lesion area

Lesion area at the focal plane was determined by focusing the laser through a coverslip (22×40 mm rectangular #1.5 (0.17 mm) thickness) onto a layer of black ink (Sharpie permanent marker, ink side was facing up and away from objective lens). Bright field mages were acquired (acquisition software: MicroManager v.2.0, camera Flash4.0 Hamamatsu, Japan) after lesion at the focal plane, as well as 2 μm above and below. Largest diameter of damage area was quantified from 5 images using ImageJ (FIJI distribution v.1.52).

### Bundle ablation

Bundle ablation was performed in adult animals that express green fluorescent protein (GFP) either pan-neuronally (strain NW1229) or in the amphid neurons (strain NG3416), obtained from *C. elegans* Genetics Center and Gian Garriaga, respectively. In all cases, an area of the bundle that included multiple neurites was brought into focus and the laser was aimed at a neurite in the bundle. Images were taken before and after lesion with MicroManager ^18^ or Nikon Elements.

### Laser axotomy and regeneration measurements

For laser microsurgery and time-lapse microscopy, *C. elegans* hermaphrodites at the fourth larval stage (L4) were mounted by placing them in a drop of cold, liquid 36% Pluronic F-127 (Sigma-Aldrich) with 1 mM levamisole (Sigma-Aldrich) solution, and pressed between two coverslips^19^. The slides were brought to room temperature, to solidify the Pluronic F-127 gel and immobilize the animals. Laser axotomy was performed using both the Yb-fibre and the nanosecond pulse laser system installed on the same microscope (ASI RAMM open frame with epifluorescence and bright field Olympus optics), for adequate comparison. In both cases the beam was focused to a diffraction-limited spot that was first located on the live image by lesion of a surface of black ink on a coverslip as described above. The targeted neuron was visually inspected immediately following laser exposure (100-500 ms) to confirm successful axotomy. In some cases, multiple laser exposures were necessary to generate a break in the nerve fibre. Axotomy of D-type motor neurons were done by severing the anterior ventral-dorsal commissures 40-50 μm away from the ventral nerve cord. Neuronal regeneration was assessed 24 hours after axotomy on the same microscope and imaging system.

### Quantification and analysis

Z-stacks were acquired both before and immediately after injury as well as 24 h post injury. Maximum intensity projections were constructed and post injury images (1 min post injury) were analysed to quantify the damage area. Images taken 24 h post injury were analysed to quantify outgrowth. Outgrowth was considered positive when a new branch was observed to be extending from the injury site. Outgrowth was quantified by measuring the length of the regenerated neurite starting from the injury site. When the neurite grew in a branched manner the distances were added. Image measurements and analysis was carried out with ImageJ software v.1.52, and statistical analysis was done with GraphPad Prism v.8.0.50.

### Statistics and interpretations of results

Most of the D motoneuron regeneration and ectopic outgrowth data is binary: we score whether or not neurons regrow or outgrow. We calculated p-values for these data by Fisher’s exact test. For regenerated length measurements and the size of the injury, we calculated p-values by the unpaired, unequal variance, two-tailed t-test. For the ink damage test, we conducted a one-way analysis of variance (ANOVA). We performed post-hoc comparisons using the Tukey test. Data are represented as average ± standard deviation (SD). * and ** indicate values that differ at p < 0.05 and 0.001 levels, respectively.

The datasets generated during and/or analysed during the current study are available from the corresponding author on reasonable request.

## Acknowledgments

We thank Chris McRaven, and Adam Perry, of Menlo Systems for support in setting up and adapting the BlueCut system. The system was purchased with funds from GH start-up resources provided by the New Jersey Institute of Technology. We also thank Michael Bastiani for advice on nanosecond pulse laser integration. Some strains were provided by the CGC, which is funded by NIH Office of Research Infrastructure Programs (P40 OD010440). This study was partially funded by grant CSCR14ERG002 from the State of New Jersey Commission on Spinal Cord Research.

## Author information

### Contributions

Approach conception: GH; ink layer experiment: MBH, VM, and SHC; amphid bundle ablation: MBH and SHC; axon regeneration experiments: MBH and VM; laser maintenance: CVG; supervision: CVG, SHC, and GH. GH wrote the first manuscript draft; all authors subsequently took part in the revision process and approve the final manuscript.

## Ethics declarations

### Competing interests

The authors declare no competing interests.

